# NINJ1 is activated by calcium-driven plasma membrane lipid scrambling during lytic cell death

**DOI:** 10.1101/2024.10.23.619800

**Authors:** Jazlyn P. Borges, Yongqian Wang, Liron David, Allen Volchuk, Brunna Martins, Ruiqi Cai, Hao Wu, Spencer A. Freeman, Neil M. Goldenberg, Michael W. Salter, Benjamin E. Steinberg

**Author notes:** These authors contributed equally to this work (Jazlyn P. Borges, Yongqian Wang).

## Abstract

NINJ1 is the terminal executioner of cellular rupture in multiple lytic cell death pathways through its clustering in the plasma membrane. Its activation trigger, however, remains unknown. We found that NINJ1-mediated plasma membrane rupture depends on calcium influx into the cell, which suffices to induce NINJ1-mediated rupture. Using genetic and pharmacologic approaches in macrophages, we show calcium drives membrane rupture through phospholipid scrambling by the calcium-activated scramblase TMEM16F. We next tested whether this calcium-activated NINJ1 mechanism is the elusive pathway by which extracellular ATP stimulates cellular rupture. We show that ATP-stimulation of P2X7R induces NINJ1-mediated cell lysis via calcium influx and TMEM16F lipid scrambling, independently of inflammasomes, pannexins and gasdermin D. Our work reveals the mechanism of NINJ1 activation and solves the long-standing mystery of ATP-induced cytolysis.

**Summary:** Elevated cytosolic calcium drives NINJ1-mediated cellular rupture during lytic cell death through plasma membrane lipid scrambling.

## Introduction

Plasma membrane rupture is the final step of lytic cell death. The release of intracellular contents into the extracellular space propagates inflammatory responses, incites further cellular death, and mediates disease (*1–3*). The transmembrane protein ninjurin-1 (NINJ1) executes terminal cellular rupture in lytic cell death through its clustering within the plasma membrane (*4*). The mechanism for triggering NINJ1 activation, however, is unknown. Given that NINJ1 is a unifying component in multiple regulated lytic cell death pathways, we hypothesized that there is a common NINJ1 triggering mechanism and that this is mediated by calcium entry into dying cells.

Increases in cytosolic calcium are observed in cell death pathways in which NINJ1 is implicated, including pyroptosis (*1*), pore-forming toxin-induced necrosis (*2*), secondary necrosis (*3*, *4*), and ferroptosis (*5*, *6*). For example, in pyroptosis, activation of the NLRP3 inflammasome leads to the formation of gasdermin D (GSDMD) pores that mediate not only the release of mature cytokines, but the influx of calcium into the cytosol (*1*, *7–9*). Calcium entry similarly occurs through pore-forming toxins, such as pneumolysin, which ultimately induce cell necrosis and NINJ1-mediated cytolysis (*2*). We therefore examined these two forms of NINJ1-mediated cell death to delineate the common mechanism by which NINJ1 is triggered to rupture the plasma membrane.

A cytolytic cell death pathway that remains enigmatic is that induced by extracellular ATP (*10*, *11*). The P2X7 receptor (P2X7R) is a proinflammatory extracellular ATP sensor that can amplify and propagate the inflammatory consequences of cell death in disease pathology (*12*, *13*). High concentrations or prolonged exposure of P2X7Rs to ATP trigger plasma membrane permeability to large molecules, referred to as the formation of a nonselective “macropore” (*10*, *11*, *14*). The notion of a P2X7R-associated macropore has been observed across a variety of cell types and experimental systems, but its molecular basis is unclear. Here, we find that cytolytic cell death induced by P2X7Rs is mediated by NINJ1 clustering, which forms the elusive macropore. Moreover, the calcium pathway we identify for pyroptosis and pore-forming toxin-induced cell death mediates P2X7R cytolysis. Thus, calcium entry is the common point of convergence for multiple cell death pathways.

## Results

### Calcium is necessary for NINJ1 aggregation and NINJ1-dependent plasma membrane rupture during pyroptosis and pore-forming toxin necrosis

To test whether calcium is necessary for NINJ1 aggregation and plasma membrane rupture in response to pyroptosis stimulation, we stimulated LPS-primed primary mouse bone marrow-derived macrophages (BMDM) to undergo pyroptosis using nigericin, a canonical NLRP3 inflammasome activator (*15*, *16*) in a complete medium (DMEM) or in medium without calcium or magnesium (PBS). When assessed in DMEM, NINJ1 migration in blue native (BN)-PAGE – which preserve native protein complexes (*17*)– basal NINJ1 migrated at approximately 40 kDa (**Fig. 1A**), consistent with previous studies (*18*, *19*). Following nigericin treatment, NINJ1shifted to a high-order molecular weight aggregate, indicative of clustering. This shift was blocked when cells were placed in PBS lacking calcium and magnesium (**Fig. 1A**). Addition of calcium to the PBS rescued NINJ1 aggregation in response to nigericin (**Fig. 1A**), indicating that extracellular calcium is necessary for NINJ1 aggregation during pyroptosis. In a complementary experiment, chelating extracellular calcium by adding EGTA to DMEM (**Fig. 1B**) suppressed LDH release from LPS-primed BMDM in response to nigericin stimulation, consistent with calcium being indispensable for plasma membrane rupture in pyroptosis. The requirement of extracellular calcium for NINJ1 clustering was not unique to pyroptosis as we obtained similar findings in macrophages stimulated to undergo necrosis due to the pore-forming toxin pneumolysin (**Supplemental Fig. S1**), which also permits calcium entry (*2*, *20*).

**Figure 1.**
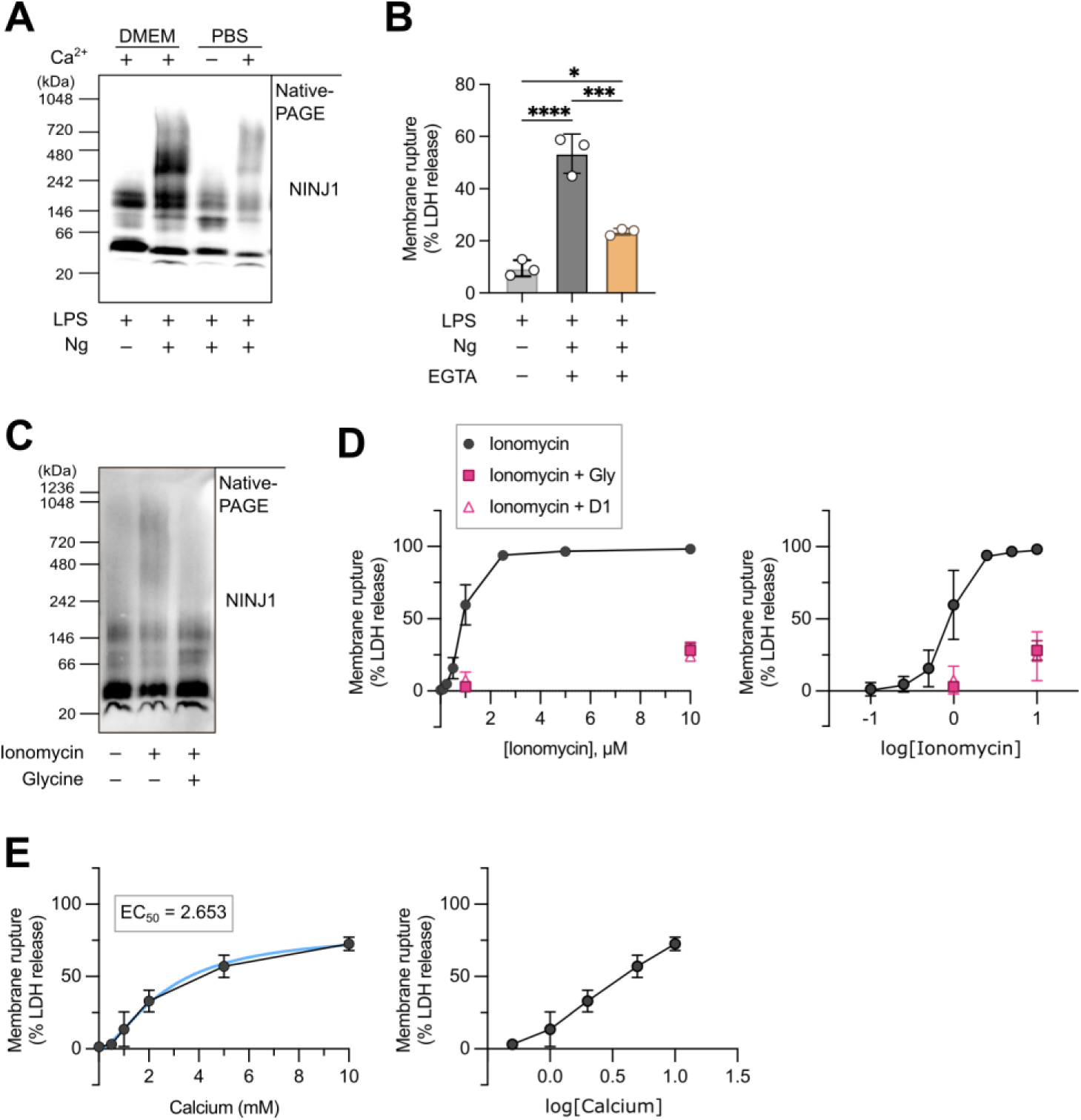
Calcium induces and is indispensable for NINJ1 aggregation and NINJ1-dependent plasma membrane rupture. **A,** BN-PAGE analysis of LPS-primed primary mouse BMDM stimulated with nigericin (20μM, 30 min) in complete media (DMEM), PBS without calcium or magnesium (−/−), or PBS(−/−) supplemented with 1.8 mM CaCl_2_. **B**, LDH release from LPS-primed BMDM stimulated with nigericin (20μM, 30 min) in DMEM complete media with or without EGTA (3.6 mM). **C,** BN-PAGE analysis of LPS-primed BMDM stimulated with nigericin (20μM, 30 min) or ionomycin (1 µM, 15 min) with or without glycine (15 mM) pre-treatment. **D,** LDH release from wildtype BMDM treated with increasing concentrations of ionomycin with or without 15 min pre-treatment with glycine (15 mM) or anti-NINJ1 (clone D1, 10 µg/mL) as indicated in the figure. The x-axis in each graph indicates the dose of ionomycin used to scale (left) or logarithmic scale (right). **E**, LDH release from primary mouse BMDM treated with ionomycin (4 µM, 15 min) in varying concentrations of extracellular calcium. The x-axis in each graph indicates the dose of ionomycin used to scale (left) or logarithmic scale (right). Data were fitted using a variable slope model to determine the EC50 of 2.7 mM (95% confidence interval 1.8-6.7 mM). The calculated fit is shown by the blue line.

### Calcium influx induces NINJ1 aggregation and NINJ1-dependent plasma membrane rupture

Having demonstrated that extracellular calcium is required for NINJ1 aggregation, we next asked whether raising intracellular calcium suffices to trigger NINJ1 aggregation and NINJ1-dependent plasma membrane rupture. To test this, we stimulated primary mouse BMDM with the calcium ionophore ionomycin. By BN-PAGE, treatment with ionomycin led to a shift in the molecular weight of NINJ1 to high-order aggregates (**Fig. 1C**) and release of LDH in a dose-dependent fashion (**Fig. 1D**). We previously showed that the cytoprotectant glycine targets NINJ1-mediated plasma membrane rupture at the level of NINJ1 clustering (*18*). Co-treatment with either glycine or with NINJ1-neutralizing antibody clone D1 (*21*) inhibits NINJ1-dependent release of LDH induced by ionomycin. These results were not specific to ionomycin, as treatment with A23187, another calcium ionophore, similarly led to dose-dependent release of LDH, which was abrogated in *Ninj1^-/-^* macrophages (**Supplemental Fig. S2**). We treated primary mouse BMDM with ionomycin in varying concentrations of extracellular calcium to determine an EC50 for NINJ1 activation of 2.7 mM (95% confidence interval 1.8-6.7 mM) as assayed by LDH release (**Fig. 1E**). From these findings, we conclude that calcium entry from the extracellular space is indispensable and sufficient to trigger NINJ1 aggregation and subsequent NINJ1-mediated plasma membrane rupture.

### Calcium induces NINJ1 activation via plasma membrane lipid scrambling by TMEM16F

A hallmark of cell death is the loss of plasma membrane lipid asymmetry through lipid scrambling (*22*, *23*), including cell death pathways in which NINJ1 clustering is triggered, such as pyroptosis (*24*), apoptosis (*25*), ferroptosis (*26*), and necroptosis (*27*). Calcium influx is known to activate lipid scrambling at the plasma membrane, in part through calcium-responsive scramblases. Therefore, we next turned our attention to whether elevated intracellular calcium could indirectly trigger NINJ1 clustering and activation by dissipating the asymmetric distribution of lipids between the exofacial and cytofacial aspects of the bilayer.

TMEM16F (gene name *Ano6*), a calcium-activated phospholipid scramblase localized at the plasma membrane (*28*) that scrambles phospholipids along their concentration gradient, has been functionally linked to lytic cell death (*26*, *29*, *30*). To test the potential requirement for TMEM16F, we generated *Ano6* knockout (*Ano6*^−/−^) in the RAW 264.7 mouse macrophage cell line. *Ano6* knockout was confirmed by western blot (**Fig. 2A**). We confirmed the *Ano6* knockout functional phenotype by labelling with annexin V, which binds exofacial phosphatidylserine (PS). Wildtype but not knockout cells exposed PS on the exofacial leaflet in response to ionomycin treatment (**Fig. 2B, Supplemental Fig. S3**). To determine whether TMEM16F is necessary for calcium-induced NINJ1-dependent lytic cell death, we stimulated wildtype or *Ano6*^−/−^ macrophages with ionomycin in the presence or absence of 15 mM glycine or clone D1 antibodies (**Fig. 2, Supplemental Fig. S3**). Like primary mouse BMDM, treatment with ionomycin ruptured wildtype RAW 264.7 cells, as evinced by LDH release into the supernatant (**Fig. 2C**). Genetic deletion of *Ano6* prevented ionomycin-induced LDH release (**Fig. 2C**). Neither glycine treatment nor NINJ1 inhibition with clone D1 offered *Ano6*^−/−^ macrophages additional protection against ionomycin-induced LDH release (**Fig. 2C**). These findings indicate that TMEM16F is necessary for calcium-induced NINJ1-dependent plasma membrane rupture in macrophages.

**Figure 2.**
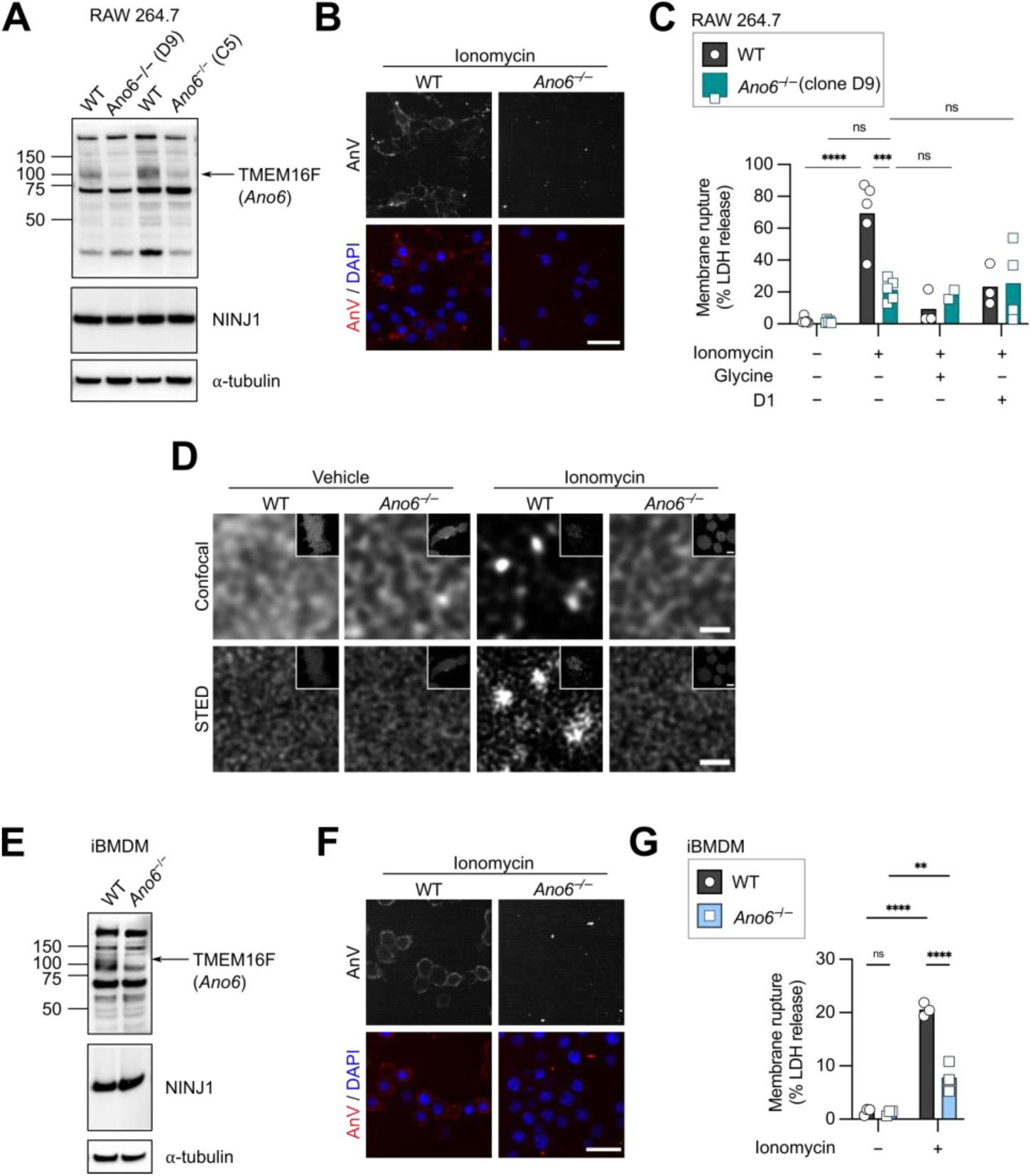
The plasma membrane calcium-dependent phospholipid scramblase TMEM16F is required for NINJ1 clustering and NINJ1-dependent rupture. **A**, Western blot of TMEM16F and NINJ1 in wildtype (WT) and *Ano6* knockout (*Ano6*^−/−^) RAW264.7 macrophages. Two clones (D9 and C5) of the *Ano6*^−/−^ were generated. α-Tubulin is shown as a loading control. **B-C,** Wildtype or *Ano6*^−/−^ RAW 264.7 cells (clone D9) treated with ionomycin (1 µM, 15 min) with or without 10-minute pretreatment with glycine (15 mM) or anti-NINJ1 antibody clone D1 (10 ug/mL) as indicated in the figure. **B,** Ionomycin treatment stimulates phosphatidylserine (PS) scrambling as assayed by Annexin V staining in the WT but not the *Ano6*^−/−^ cells. Scale bar 60 µm. **C,** Ionomycin treatment stimulates cell rupture as assayed by LDH release in the WT cells, which is suppressed by genetic deletion of *Ano6.* Neither glycine nor clone D1 provide *Ano6*^−/−^ cells additional protection against ionomycin-induced LDH release. *** p < 0.001, **** p < 0.0001, ns not significant by two-way ANOVA with Tukey post-test. **D**, Representative stimulated emission depletion microscopy (STED) and corresponding confocal images of endogenous plasma membrane NINJ1 in wildtype and *Ano6*^−/−^ RAW 264.7 cells treated with ionomycin (1 µM, 15 min) or vehicle (DMSO). NINJ1 clustering in wildtype cells treated with ionomycin is prevented by *Ano6* genetic knockout. **E,** Western blot of TMEM16F and NINJ1 in wildtype (WT) and *Ano6* knockout (*Ano6*^−/−^) immortalized bone marrow-derived macrophages (iBMDM). α-Tubulin is shown as a loading control. **F-G**, Wildtype or *Ano6*^−/−^ iBMDM were treated with ionomycin for 15 minutes. **F**, Ionomycin treatment leads to LDH release from WT cells is suppressed by genetic deletion of *Ano6.* Scale bar 60 µm. **G**, Ionomycin-stimulated PS scrambling (indicated by Annexin V labelling) is prevented by genetic deletion of in *Ano6*^−/−^.

To rule out the possibility that NINJ1 distribution is altered in *Ano6*^−/−^ cells, we performed super-resolution microscopy by stimulated emission depletion (STED) comparing wildtype and knockout cells. Consistent with our previous studies in primary mouse BMDM (*18*), NINJ1 was distributed in small puncta on the surface of RAW 264.7 cells at rest, regardless of genotype (**Fig. 2D**). In wildtype cells treated with ionomycin, NINJ1 puncta appeared to increase in intensity and decrease in density, while the distribution of NINJ1 in *Ano6*^−/−^ cells remained largely unchanged compared to controls (**Fig. 2D**). To rule out a RAW 264.7 cell line-specific effect, we also generated *Ano6* knockouts in immortalized BMDM (iBMDM) (**Fig. 2E**). Like RAW 264.7 cells, *Ano6*^−/−^iBMDM did not undergo calcium-induced lipid scrambling (**Fig. 2F**) and were protected from ionomycin-induced lytic cell death (**Fig. 2G**). From these collective data, we conclude that calcium-induced NINJ1 aggregation requires TMEM16F-mediated plasma membrane lipid scrambling in macrophages.

### BzATP stimulates P2X7R-dependent cell rupture and requires NINJ1

We have demonstrated that calcium influx induces and is indispensable for NINJ1 activity through induction of plasma membrane phospholipid scrambling. We next sought to determine whether this NINJ1 activation pathway explains how extracellular ATP activates lytic cell death. We examined the possibility that P2X7R activation stimulates NINJ1 clustering by increasing calcium permeability to activate plasma membrane lipid scrambling.

Stimulation of primary mouse peritoneal macrophages with BzATP, a potent P2X7R agonist (*31*, *32*), lead to cellular rupture as evinced by release of LDH into the supernatant (**Fig. 3A**). To further characterize the cell permeabilization induced by P2X7R activation, we employed YO-PRO-1, a 630 Da monomeric cyanine nucleic acid stain that fluoresces when bound to DNA and which is commonly employed for membrane permeabilization studies (*33*). We detected two principal phases of YO-PRO-1 uptake: An initial slow linear phase starting at the time of BzATP treatment (**Fig. 3** and **Supplemental Fig. S4**), followed by a fast phase in which YO-PRO-1 fluorescence undergoes a drastic increase before plateauing after approximately 20 minutes (**Fig. 3B-C**). The kinetics of dye uptake at the individual cell level suggested a cataclysmic cellular event in which the cell becomes rapidly and grossly permeable to the YO-PRO-1 dye at a rate approximately 100-fold greater than the initial linear phase (**Fig. 3B-C**). Indeed, the fast phase of YO-PRO-1 uptake upon treatment with BzATP is consistent with the cell rupture assayed by LDH release (**Fig. 3A**). Linear phase data of YO-PRO-1 uptake are shown in **Supplemental Fig. S4** and **Table 1**.

**Figure 3.**
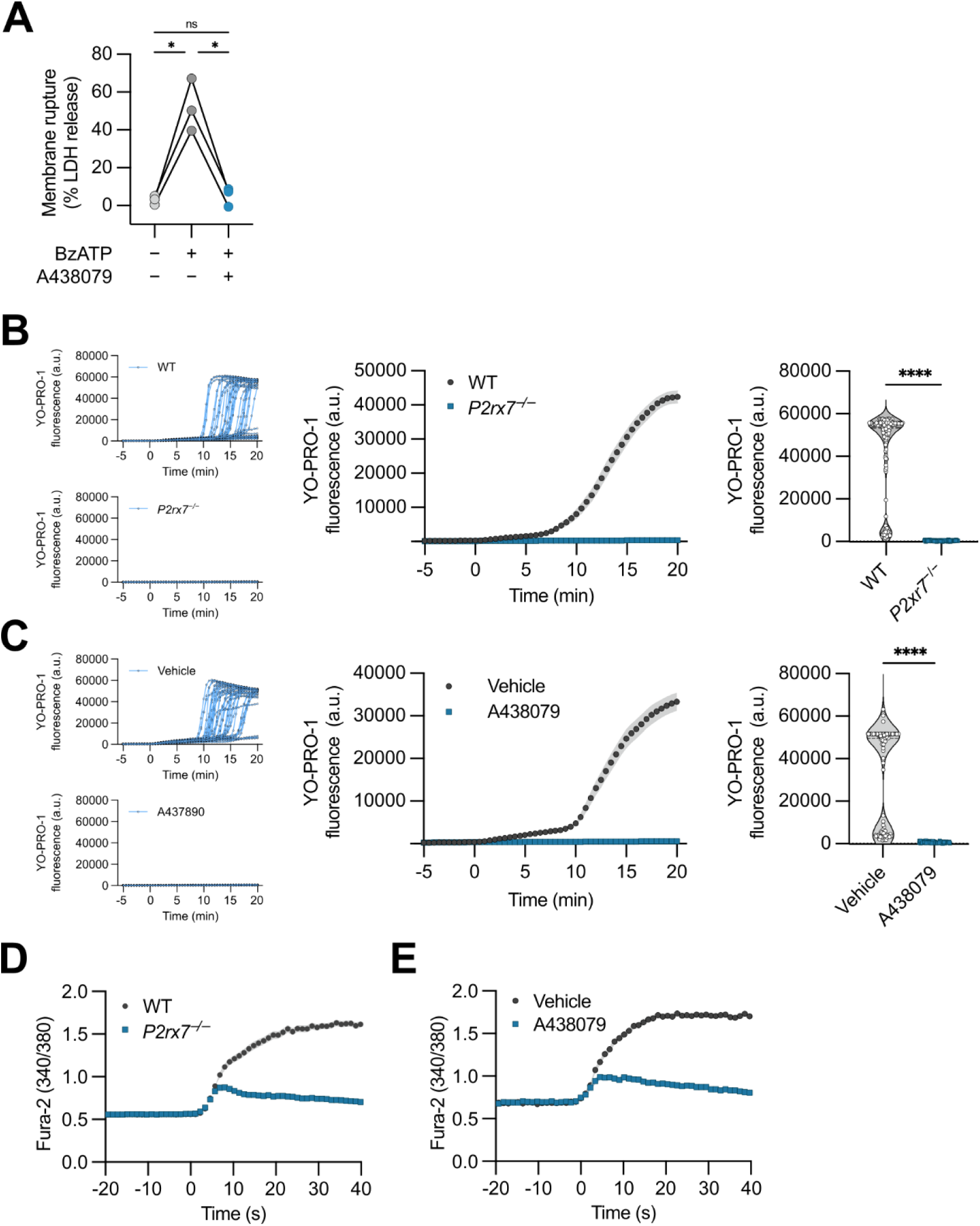
BzATP stimulates P2X7R-dependent lytic cell death. **A,** Stimulation of peritoneal macrophage with BzATP induces lytic cell death as assayed by LDH release into the supernatant. Pretreatment with A238079 prevents BzATP-induced cytotoxicity. * *p* < 0.05 by repeated measures one-way ANOVA with Geisser-Greenhouse correction and Tukey post-test from *n* = 3 independent experiments. **B-C**, YO-PRO-1 fluorescence was measured in primary mouse peritoneal macrophages at baseline, followed by stimulation with 300 µM BzATP over the course of 20 min. The leftmost graph in each panel indicates the average fluorescence intensity of YO-PRO-1 within cells over time (minutes). The middle graph depicts representative single cell tracings of YO-PRO-1 dye uptake. The rightmost graph (violin plot) in each panel indicates average YO-PRO-1 fluorescence intensity within individual cells (circles, squares) at 20 minutes following BzATP treatment. Peritoneal macrophages were stimulated with or without P2X7R perturbation by genetic knockout (*P2rx7* KO) (**B**) or pre-treatment with A438079 (**C**). **D-E**, Intracellular calcium measurements in primary mouse peritoneal macrophages before and after the addition of 300 µM BzATP. Graph indicates the ratio of fluorescence intensity of Fura-2 at 340 and 380 nm over time following BzATP treatment. Timepoints are shown relative to the first recorded measurement following BzATP treatment (t = 0 s). Peritoneal macrophages were stimulated with or without P2X7R perturbation by genetic knockout (*P2rx7* KO) (**D**) or pre-treatment with A438079 (**E**).

**Table 1.**
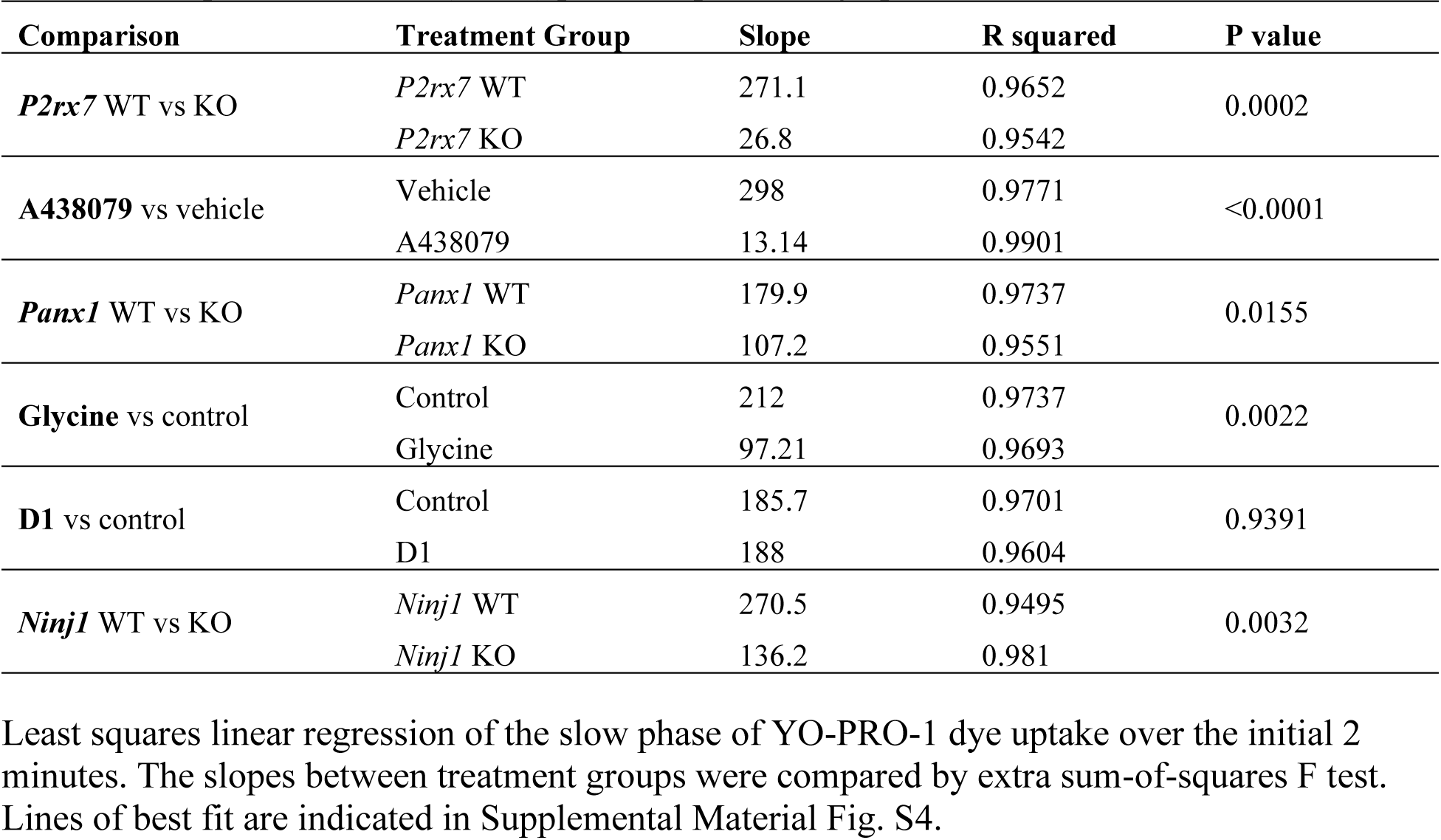
Comparison of slow (linear) phase slopes from graphed YO-PRO-1 data.

YO-PRO-1 dye uptake required P2X7R as it was absent in *P2×7r* knockout cells (**Fig. 3B**) or following pre-treatment with the P2X7R antagonist A438079 (*34*) (**Fig. 3C**). LDH release was similarly prevented by pretreatment with A438079 (**Fig. 3A**). To confirm P2X7R inhibition, we conducted single-cell calcium imaging using Fura-2, a ratiometric calcium sensor (*35*, *36*). We detected a rapid increase in intracellular calcium, which saturated within 10 seconds following BzATP treatment of wildtype macrophages (**Fig. 3D-E**). This increase was largely abolished by genetic ablation of *P2×7* (**Fig. 3D**) or by pre-treatment with A43079 (**Fig. 3E**). The small increase in intracellular calcium we observed when P2X7R was genetically deleted or pharmacologically inhibited likely represent activation of other purinergic receptors, such as the P2Y2 receptor, which are weakly activated by BzATP (*37*) and are known to release intracellular calcium stores via IP_3_ production (*38*).

Prolonged or high-dose ATP activation of P2X7R is widely known to facilitate the release of larger (> 900 Da) molecules by the formation of the P2X7R macropore (*11*). Whether NINJ1 is responsible for the P2X7R macropore activity is not known. Thus, we next asked whether the cataclysmic nonlinear phase of P2X7R-mediated YO-PRO-1 uptake reflects NINJ1-driven cell rupture. Pre-treatment of macrophages with glycine suppressed LDH (**Fig. 4A**) and prevented the fast, nonlinear phase of YO-PRO-1 uptake (**Fig. 4B**). Glycine treatment did not interfere with BzATP-induced calcium influx (**Supplemental Fig. S5**), indicating that glycine does not perturb P2X7R activation or its calcium conductance. Targeting NINJ1 with the neutralizing antibody clone D1 (*21*) similarly prevented the fast phase of YO-PRO-1 uptake (**Fig. 4C**). To complement these pharmacologic investigations, we used *Ninj1*^−/−^ immortalized bone marrow-derived macrophages (iBMDM) (*18*). *Ninj1* knockout cells maintained their capacity to respond to BzATP with an initial linear YO-PRO-1 uptake but did not demonstrate the fast secondary phase (**Fig. 4D**).

**Figure 4.**
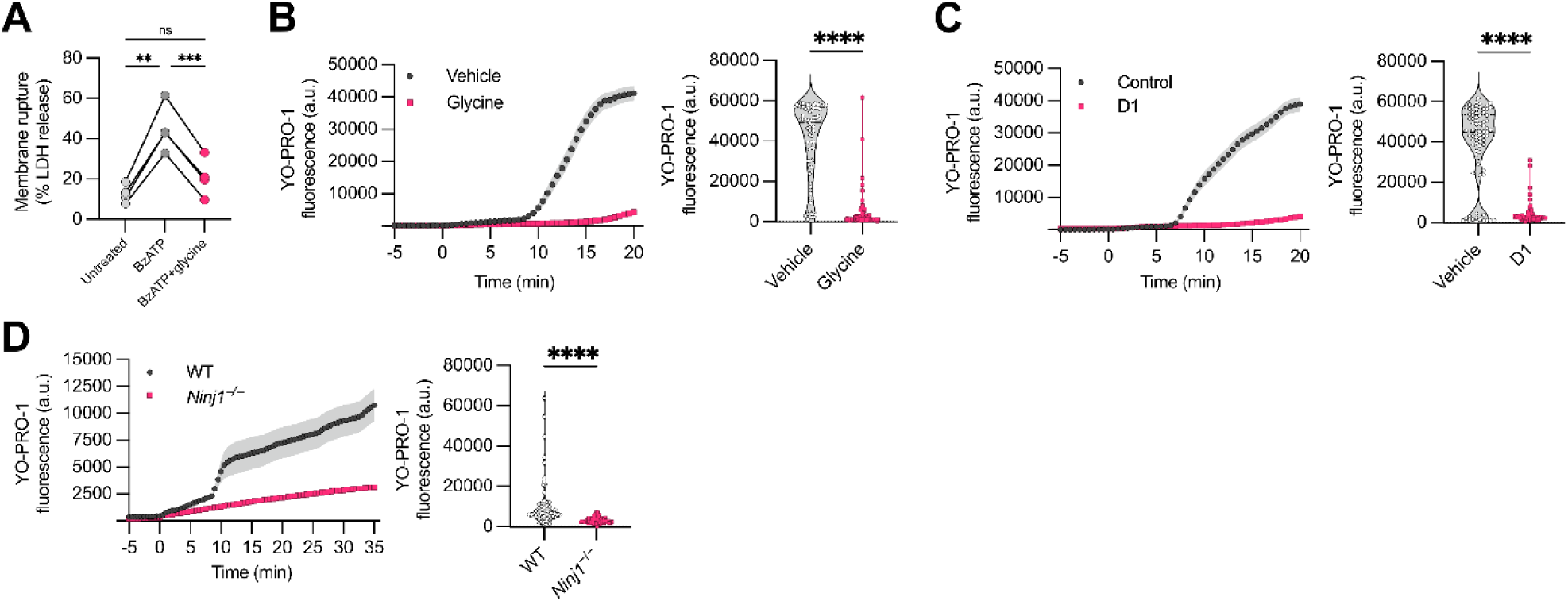
The fast phase of BzATP-stimulated YO-PRO-1 uptake requires NINJ1. YO-PRO-1 fluorescence was measured in mouse macrophages at baseline, followed by stimulation with 300 µM BzATP over the course of 20 min. The left graph in each panel indicates the average fluorescence intensity of YO-PRO-1 within cells over time (minutes). The right graph in each panel indicates average YO-PRO-1 fluorescence intensity within individual cells (circles, squares) at the endpoint of BzATP treatment. **A-B,** Wildtype primary mouse peritoneal macrophages were pre-treated for 10 minutes with glycine (15 mM) prior to stimulation with BzATP (300 µM) for 20 min. **B,** Stimulation of peritoneal macrophage with BzATP induces lytic cell death as assayed by LDH release into the supernatant, which was suppressed by pretreatment with glycine (1 mM). ** *p* < 0.01, *** *p* < 0.001 by repeated measures one-way ANOVA with Geisser-Greenhouse correction and Tukey post-test from *n* = 4 independent experiments. **C,** Wildtype primary mouse peritoneal macrophages were pre-treated for 10 minutes with anti-NINJ1 antibody (clone D1; 10 µg/mL). **D,** Wildtype or *Ninj1*^−/−^ immortalized bone marrow-derived macrophages (iBMDM) were stimulated with BzATP for a total of 35 min. **** *p* < 0.0001 by Mann-Whitney test; *n* = 62-80 cells per group from 2-3 independent experiments; primary cells derived from 2-3 animals (**A**, **C, D**).

### P2X7R-mediated cell rupture is independent of pannexin-1, NLPR3, and GSDMD

P2X7R activation has been associated with pannexin-1 and NLRP3-mediated pyroptosis (*11*, *12*, *39–42*). We next evaluated whether these intermediary proteins contribute to BzATP-induced NINJ1 activation as assayed by the second, fast non-linear phase of YO-PRO-1 dye uptake. Although pannexin-1 has been implicated in P2X7R macropore formation (*11*, *12*), we did not detect a difference in YO-PRO-1 dye uptake in *Panx1*^−/−^ macrophages relative to wildtype cells (**Fig. 5A**, **Supplemental Fig. S4**) or in cells treated with the pannexin inhibitor 10PANX (**Fig. 5B**). Given that calcium influx and P2X7R activation have been implicated in NLRP3 inflammasome activation (*39–42*), we asked whether the nonlinear phase of BzATP-stimulated YO-PRO-1 uptake represents pyroptotic cell death. To test this, we evaluated whether the second phase requires NLRP3 inflammasome activation or GSDMD. We pharmacologically inhibited the NLRP3 inflammasome with MCC950 (*43*) and evaluated the role of GSDMD by knocking out *Gsdmd* (*7*). Pre-treatment of wildtype primary mouse macrophages with MCC950 did not inhibit the BzATP-induced increase in YO-PRO-1 dye uptake at 20 min seen in vehicle control-treated cells (**Fig. 5C**). Similarly, following BzATP stimulation, YO-PRO-1 uptake in *Gsdmd*^−/−^ macrophages increased comparably to wildtype cells (**Fig. 5D**). Together, these data provide compelling evidence that P2X7R macropore formation depends on NINJ1 independent of pannexin-1, NLRP3 inflammasome activation or GSDMD.

**Figure 5.**
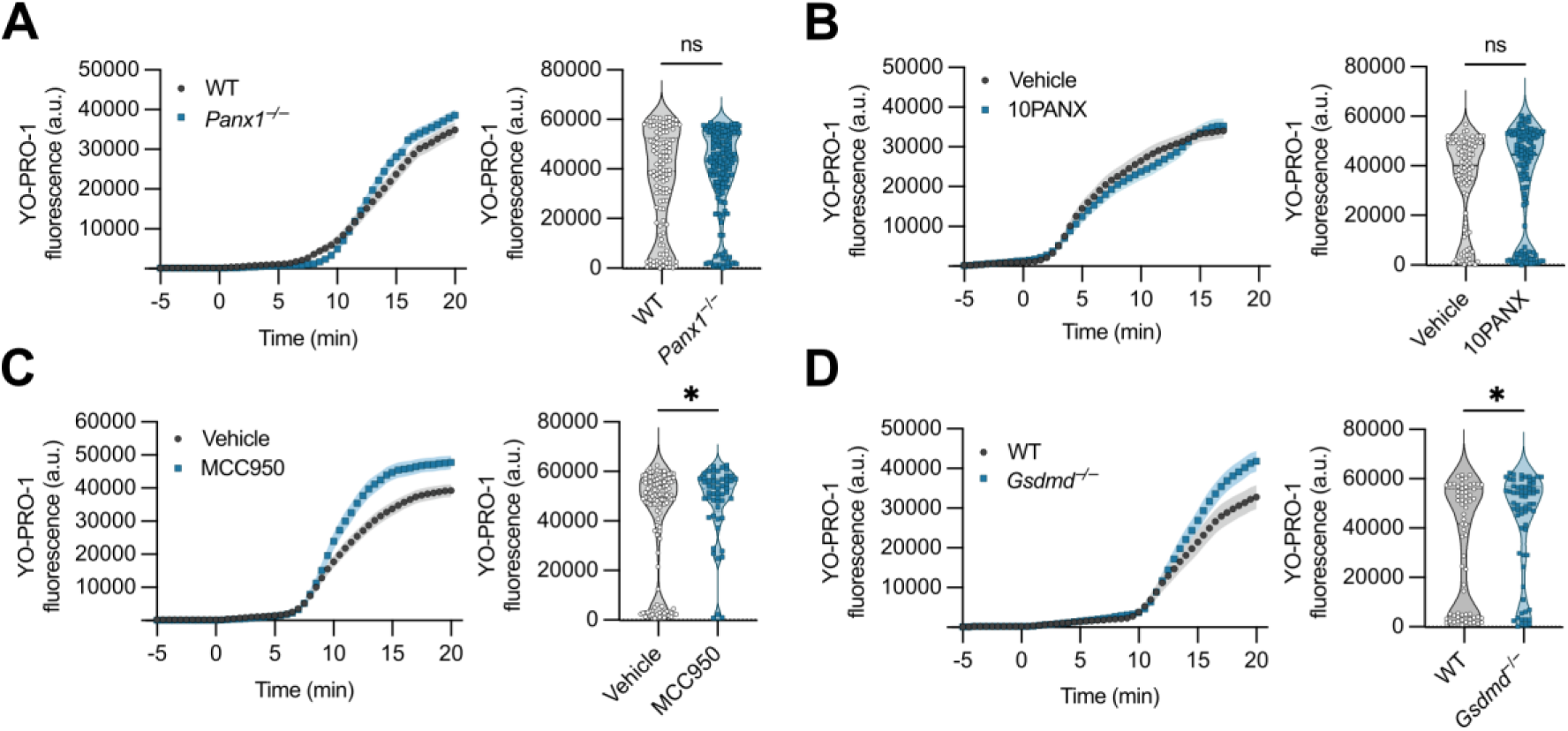
BzATP-stimulated lytic cell death is independent of pannexin-1, NLRP3, and GSDMD. YO-PRO-1 fluorescence was measured in primary mouse peritoneal macrophages at baseline, followed by stimulation with 300 µM BzATP over the course of 20 min. **A-D**, The leftmost graph in each panel indicates the average fluorescence intensity of YO-PRO-1 within cells over time (minutes). The middle graph depicts representative single cell tracings of YO-PRO-1 dye uptake. The rightmost graph (violin plot) in each panel indicates average YO-PRO-1 fluorescence intensity within individual cells (circles, squares) at 20 minutes following BzATP treatment. Peritoneal macrophages were stimulated with or without pannexin-1 perturbation by genetic knockout (*Panx1* KO) (**A**) or pharmacologic inhibition (10PANX) (**B**), NLRP3 inflammasome inhibition by pre-treatment with MCC950 (**C**), or gasdermin D perturbation by genetic knockout (*Gsdmd* KO) (**D**). * *p* < 0.05, **** *p* < 0.0001 by Mann-Whitney test; *n* = 53-160 cells per group from 2-3 animals.

### P2X7R activation stimulates NINJ1 aggregation and fast YO-PRO-1 uptake in a calcium-dependent manner

To evaluate the mechanism underlying NINJ1 activation by BzATP, we again used BN-PAGE. Basal NINJ1 migrated at approximately 40 kDa, whereas NINJ1 migration shifted to high-order molecular weight aggregates following BzATP treatment (**Fig. 6A**). P2X7R-mediated lytic cell death has been associated with calcium influx and downstream calcium-responsive signaling (*11*), which is consistent with our model of NINJ1 activation. Placing cells in a calcium-free medium prevented NINJ1 aggregation in response to BzATP (**Fig. 6A**), suggesting that P2X7R-mediated NINJ1 aggregation depends on the influx of extracellular calcium. We next aimed to confirm whether calcium is necessary for the fast nonlinear phase of YO-PRO-1 uptake. Chelation of calcium with EGTA (**Fig. 6B**) prevented the fast YO-PRO-1 uptake. Extracellular calcium influx therefore facilitates P2X7R-dependent NINJ1 aggregation and NINJ1-mediated cell lysis.

**Figure 6.**
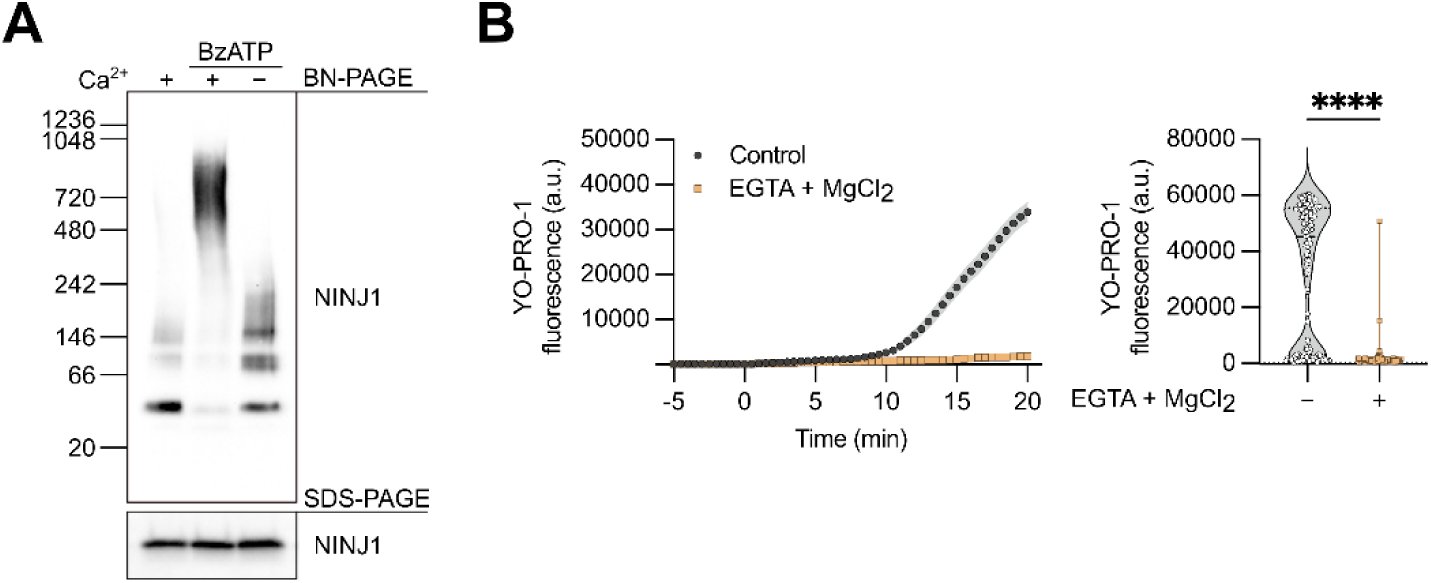
BzATP stimulates calcium-dependent NINJ1 aggregation and YO-PRO-1 uptake. **A,** BN-PAGE analysis of primary mouse peritoneal BMDM left untreated (lane 1) or stimulated with BzATP (30 μM, 20 min) in ECS with (lane 2) or without calcium (lane 3). **B,** YO-PRO-1 fluorescence was measured in primary mouse peritoneal macrophages at baseline, followed by stimulation with 300 µM BzATP over the course of 20 min in ECS with or without 1.3 mM EGTA. The leftmost graph in each panel indicates the average fluorescence intensity of YO-PRO-1 within cells over time (minutes). The rightmost graph in each panel indicates average YO-PRO-1 fluorescence intensity within individual cells (circles, squares, or triangles) at the endpoint of BzATP treatment. **** *p* < 0.0001, ns not significant by Mann-Whitney test; *n* = 72-134 cells per group from 2-3 animals.

## Discussion

We found that calcium is necessary for NINJ1 aggregation during pyroptosis and necrosis, and that treatment with calcium ionophores stimulates NINJ1 aggregation and NINJ1-dependent plasma membrane rupture in a TMEM16F-dependent fashion. Our data therefore define sustained intracellular calcium increase as the first characterized trigger of NINJ1 activation. Elevated intracellular calcium is a common feature across pyroptosis (*1*, *44*, *45*) and pore-forming toxin-induced necrosis (*2*), consistent with our conclusions. Our findings are also consistent with a recent study by Ramos *et al*, which determined that, in ferroptosis, calcium influx precedes and is independent of NINJ1 activation (*46*). Lastly, although NINJ1 is dispensable for plasma membrane rupture in response to necroptosis, we have previously demonstrated that NINJ1 aggregates in response to necroptotic stimuli (*18*). In that case, we posit that calcium influx mediated by MLKL, the pore-forming protein responsible for necroptosis, facilitates NINJ1 aggregation via calcium influx and plasma membrane lipid scrambling.

Cell swelling has been proposed as a trigger of NINJ1 activation (*47*, *48*). In this regard, it is noteworthy that hypotonic cell swelling (*49*) has been shown to stimulate calcium influx via mechanosensitive ion channels such as TRPV4 (*50*) or PIEZO1 (*6*). For example, hypotonic shock induces calcium influx in RAW 264.7 cells via PIEZO1 (*6*). Additionally, cell volume changes induce calcium-dependent PS exposure (*51*), which has been proposed to occur via TMEM16F. In this way, our findings are consistent with the study by Dondelinger *et al,* which indicates that hypotonic shock leads to NINJ1 activation (*48*).

Our studies specifically implicate TMEM16F, the most calcium-responsive lipid scramblase in macrophages (*29*), in the terminal events of cellular rupture. We observed that knocking out *Ano6* prevents NINJ1 aggregation and membrane rupture, and that *Ano6* KO minimize scrambling. Following its calcium-dependent activation, TMEM16F mediates rapid phospholipid scrambling (*28*), however, we have not tested whether blocking scrambling *per se* prevents NINJ1 aggregation and cell rupture. These may ultimately represent epiphenomena of another and that TMEM16F is serving an alternative function in NINJ1-mediated cell death.

Plasma membrane lipid scrambling alters the local distribution of varying lipid species and the biophysical properties of the membrane. Both may contribute to NINJ1 activation. Phosphatidylserine and phosphatidylinositol-4-phosphate have been implicated in the formation of NINJ1 aggregates and therefore NINJ1-mediated plasma membrane rupture (*52*, *53*). Plasma membrane lipid scrambling may therefore mobilize important lipid species, such as phosphatidylserine, that associate with NINJ1 to facilitate its clustering. Alternatively, lipid scrambling may cause large-scale biophysical changes of the plasma membrane, such as alterations in lipid packing, membrane electrostatic properties, or the formation of local non-bilayer phases (*54*). These disruptions to lipid asymmetry impact membrane curvature, fluidity, and tension as well as protein conformation and their association with the plasma membrane (*55*). Thus, specific lipid species or altered biophysical membrane properties may be licensing NINJ1 clustering, which remain to be empirically determined.

While our studies have focused on TMEM16F given its prominent role in macrophages, other lipid scramblases may contribute to NINJ1 activation in different cell types and death pathways. For example, the caspase-activated scramblase Xkr8 is necessary for lipid scrambling following apoptosis stimulation, while TMEM16F is dispensable (*56*). Additionally, the scramblase XK, which complexes with the lipid transporter VPS13A, has been proposed to be important for P2X7R-mediated phospholipid scrambling and ATP-induced propidium iodide uptake as measure of cell membrane permeability in a TMEM16F-deficient cell line and primary mouse T cells (*57*). Other scramblases may compensate in the context of TMEM16F deficiency in a cell type-dependent fashion Notably, TMEM16F has also been associated with plasma membrane repair processes in lytic cell death pathways (*58–61*). For example, the ESCRT-III machinery excises GSDMD-containing vesicles to enable membrane repair during pyroptosis (*1*) and MLKL-containing vesicles during necroptosis (*61*). ESCRT-III and other membrane repair mechanisms may initially counteract TMEM16F-dependent NINJ1 clustering; however, is also possible that TMEM16F is involved both in membrane repair processes and in NINJ1 clustering. Given that ESCRT-III-mediated membrane repair is an ATP-dependent process (*62*), and NINJ1-dependent plasma membrane rupture occurs after the loss of mitochondrial membrane potential and ATP depletion (*47*, *63*), it is plausible that in the absence of available repair mechanisms, cells succumb to NINJ1-dependent plasma membrane rupture following phospholipid scrambling.

The nature of the macropore responsible for P2X7R-mediated lytic cell death has been a controversial topic of investigation and has evaded identification (*11*). Here, we position NINJ1 clusters as the effective P2X7R macropore that facilitates ATP-stimulated lytic cell death independently of pannexin-1, NLRP3, and GSDMD. Although P2X7R-dependent lytic cell death has been linked to NLRP3 inflammasome activation and pyroptosis, priming cells with LPS to induce expression of inflammasome machinery may bias cells towards NLRP3-dependent pyroptosis. Indeed, previous studies have shown that priming enhances P2X7-mediated IL-1β and LDH release in peritoneal mouse macrophages (*64*), suggesting that LPS licenses P2X7R-dependent NLRP3 inflammasome activation and GSDMD-mediated pyroptosis in this context. NINJ1 likely serves as the terminal effector of cell rupture in GSDMD-dependent and independent instances.

In summary, we discovered that pyroptosis stimulation or ATP-mediated P2X7R activation leads to NINJ1 clustering and plasma membrane rupture in a calcium influx- and TMEM16F-dependent fashion. Moreover, our work revealed that NINJ1 mediates the cytolysis attributed to the elusive P2X7R macropore. It will be important for future studies to delineate how NINJ1 senses calcium-induced plasmalemmal lipid scrambling during cell death to cluster and ultimately rupture the cell.

## Materials and Methods

### Animals and primary cell culture

#### Animals

Wild-type C57BL/6 animals were purchased from Jackson Laboratories (strain #000664). *Gsdmd*^−/−^ and *Ninj1*^−/−^ animals were a kind gift from Dr. Nobuhiko Kayagaki and Dr. Vishva Dixit (Genentech Inc.) (*7*, *19*). The *P2rx7* null mouse line was generated by crossing *P2rx7 fl/fl* mice (TCP, req. 54620018) with CMV-Cre mice (JAX, Strain #:006054), while the *Panx1* null mouse line was obtained from Dr. Peter Carlen’s lab (University Health Network). All animal procedures were conducted under protocols approved by the Animal Care Committee at The Hospital for Sick Children and in accordance with animal care regulation and policies of the Canadian Council on Animal Care. Mice were housed in same-sex polycarbonate cages with *ad libitum* access to food and water. Housing rooms were temperature and humidity controlled with 14:10 h light:dark cycles.

#### Bone marrow-derived macrophages

Primary bone marrow-derived macrophages (BMDM) were harvested from the femurs of mixed-sex cohorts of wild-type mice. The bones were cleaned, the ends cut and centrifuged to collect bone marrow into sterile PBS. Following a wash in PBS, the cells were plated in DMEM (Wisent Bioproducts) supplemented with 10% fetal bovine serum (R&D Systems, S12450), 100 U/ml penicillin, and 100 μg/ml streptomycin (Wisent Bioproducts, 450-201), and 10 ng mL^-1^ M-CSF (315-02; Peprotech Inc, Cranbury, NJ) and maintained at 37°C and 5% CO_2_. After 5 days of culture, the BMDM were detached from the dishes in TBS with 5 mM EDTA, resuspended in fresh medium and plated.

#### Peritoneal macrophages

Peritoneal macrophages were obtained through peritoneal lavage of adult mice between 8-12 weeks of age using cold PBS (Wisent, 311-012-CL). These macrophages were subsequently enriched and cultured at 37°C with 5% CO2 in RPMI-1640 media (Thermo Fisher, A1049101) supplemented with 10% fetal bovine serum and 1% penicillin and streptomycin. Prior to experimentation, cells were cultured overnight on poly-D-lysine-coated 35mm glass-bottom dishes (MatTek, P35GC-1.0-14-C).

### Knockout cell lines

iBMDM and RAW cells were transfected with custom CRISPR gRNA plasmid DNA (U6-gRNA:CMV-Cas-9–2 A-tGFP; Sigma-Aldrich, Oakville, Canada) using FuGENE HD transfection reagent (Promega, Madison, WI, USA) as previously described (*18*). The *Ninj1* target region sequence was GCCAACAAGAAGAGCGCTG and the *Ano6* target region sequence was CTTCTCGTAGATCGTGTTG. 24 hr later, the cells were FACS sorted for GFP into 96 well plates. Individual colonies were expanded and tested for NINJ1 or TMEM16F protein expression by western blot. Primary antibody against TMEM16F was used at 1:1000 (Sigma Aldrich, HPA038958). Wildtype, *Ano6* knockout, and *Ninj1* knockout macrophages were maintained in DMEM supplemented with 5% FBS (RAW cells) or 10% FBS (iBMDM) and 100 U/ml penicillin and 100 μg/ml streptomycin at 37 °C and 5% CO_2_.

### Cell stimulation

Pyroptosis was activated in primary mouse BMDM primed with (0.5 µg/ml) LPS from *E. coli* serotype 055:B5, which was first reconstituted at a stock concentration of 1 mg/ml. Cells were primed with LPS for a total of 5 hours, during which pyroptosis was induced with nigericin (Sigma N7143; stock 10 mM in ethanol) for the final 30 min at a concentration of 20 µM. Ionomycin (Sigma, 407950) and A23187 (Sigma, C7522) were reconstituted in stock solutions at 10 mM in DMSO. Cells were stimulated with ionomycin as indicated in the figure legends or 10 µM A23187 in serum-free DMEM for 15 or 30 min at 37°C, or otherwise as indicated. Anti-NINJ1 neutralizing antibody clone D1 (generous gift from Dr. Nobuhiko Kayagaki and Dr. Vishva Dixit, Genentech) was used to pre-treat cells at 10 µg/mL for 15 min prior to stimulation (*21*). Cells were pre-treated with glycine (Sigma, G8898) at 15 mM or otherwise as indicated in the figure legends.

### YO-PRO dye uptake assays

For the investigation of P2X7R-mediated pore formation, cells were incubated with 2.5 μM YO-PRO-1 Iodide dye (Thermo Fisher, Y3603) in an extracellular recording solution (ECS) with the following composition (in mM): NaCl 140, KCl 5.4, CaCl_2_ 1.3, HEPES 10, glucose 33, pH 7.35, and an osmolarity of 315-320 mOsm. These experiments were conducted at room temperature (22–25 °C), and all drugs were administered in ECS, along with 2.5 μM YO-PRO. To initiate P2X7R activation, cells were stimulated with 300 μM BzATP (Sigma, B6396-25MG). YO-PRO fluorescence was recorded and measured at 30-second intervals, with the recording commencing after a 5-minute pre-BzATP baseline and continuing for a total duration of 30 minutes using the PTI EasyRatioPro Ca++ Imaging System (HORIBA Scientific). In experiments examining the effects of extracellular calcium, peritoneal macrophages were washed with Calcium-Free ECS (CFECS, where CaCl_2_ was replaced with 1.3 mM MgCl_2_) before being incubated in 2.5 μM YO-PRO in CFECS, which also contained 1.3 mM EGTA. For intracellular calcium effect experiments, macrophages were washed with either ECS or CFECS, and then incubated in 2.5 μM YO-PRO in ECS or CFECS, with an additional 10 μM BAPTA-AM. For experiments involving pharmacologic inhibition of NINJ1, cells were pre-treated with 10 μg/mL anti-NINJ1 antibody (clone D1) or 15 mM glycine in ECS for 10 min prior to cell stimulation with 300 µM BzATP.

### Intracellular calcium imaging

Cells were incubated with 2.5 μM of fluorescent Ca^2+^-indicator dye Fura-2-AM (Thermo Scientific, F122) in ECS described above and supplemented with 1% BSA for 30 minutes, followed by thorough ECS rinses and a subsequent 30-minute incubation in ECS at room temperature for hydrolysis of the AM moiety. Cells were thoroughly rinsed with ECS prior to imaging recordings. Intracellular calcium dynamics were measured using the PTI EasyRatioPro Ca++ Imaging System (HORIBA Scientific), capturing emissions at 340 nm and 380 nm. Changes in intracellular calcium levels were assessed by calculating the fluorescence ratio of 340/380 after dynamic background subtraction.

### Pharmacological inhibitors for YO-PRO uptake and calcium imaging studies

For pharmacologic inhibition in YO-PRO assay and intracellular calcium measurements, cells were pretreated for 10 minutes with the indicated inhibitor before subsequent stimulations with BzATP. Inhibitors were used as follows: A438079 (100 μM; Tocris #2972) 10PANX (250 μM; Tocris #3348); MCC950 (1 μM; HelloBio #HB4636), glycine (1-15 mM as indicated in the figure legends; Sigma #G8898), EGTA (100 μM; Sigma #03777), anti-NINJ1 neutralizing antibody (clone D1; 10 µg/mL) and BAPTA-AM (10 μM, Abcam #ab120503).

### LDH assay

Cells were seeded at 300 000 cells per well in 12-well plates, treated as indicated, and cytotoxicity assayed by LDH release assay the following day. Following treatment, the cell culture supernatants were collected, cleared of debris by centrifugation for 5 min at 500 x *g*. The cells were lysed in lysis buffer provided in the LDH assay kit (Invitrogen, C20300). Supernatants and lysates were assayed for LDH using an LDH colorimetric assay kit as per the manufacturer’s instructions (Invitrogen, C20300).

### Blue Native-PAGE, SDS-PAGE, and western blotting

Mouse macrophages were lysed with native-PAGE lysis buffer (150 mM NaCl, 1% Digitonin, 50 mM Tris pH7.5, and 1× Complete Protease Inhibitor). Following centrifugation at 20,800× *g* for 30 min, lysates were mixed with 4× NativePAGE sample buffer and Coomassie G-250 (ThermoFisher) and resolved using NativePAGE 3–12% gels. For SDS-PAGE, cells were washed with 1× PBS and lysed in RIPA lysis buffer containing protease inhibitors (Protease inhibitor tablet, Pierce A32955). Proteins were resolved using NuPAGE 4–12% Bis-Tris gels (Invitrogen), transferred to 0.2 µm PVDF (polyvinylidene difluoride) membranes, and immunoblotted using a monoclonal rabbit anti-mouse NINJ1 antibody (clone 25; kind gift from Dr. Kayagaki and Dr, Dixit at Genentech, Inc.) (*19*).

### Immunoprecipitation

Immuno-precipitation experiments were conducted for the following proteins: NINJ1-GFP-FLAG with P2X7-Tdtomato, NINJ1-GFP-FLAG alone and P2X7-Tdtomato alone. Proteins were co-expressed in expi293F mammalian cells. For each pull-down, 25 ml cells at 3×10^6^ cells/ml were transfected with 20 μg plasmid of each construct using lipofectamine 2000 as per the manufacturer’s instructions. Cells were harvested 24 h post-transfection and pulled down using FLAG beads for pulling down NINJ1-FLAG. Western blot analysis was performed for each IP experiment with the following fractions – loading fraction, elution and beads. In addition, as a control we performed FLAG pulldowns for only NINJ1-GFP-FLAG, P2X7-Tdtomato. Detection of NINJ1-GFP-FLAG was utilized by using human NINJ1 antibody (1:1000. R&D-AF5105) and P2X7 antibody (1:1000, Novus-NBP1-20180).

### Super-resolution microscopy

#### Sample preparation

Wildtype and *Ano6*^−/−^ RAW 264.7 macrophages were cultured on 1.5H glass coverslips. Cells were treated as indicated in the figure legend, then washed once with PBS and fixed in 4% paraformaldehyde in PBS at room temperature for 15 min. Cells were washed 3 times in PBS, then blocked in 10% donkey serum in PBS for 1 h. Cells were incubated with rabbit monoclonal anti-mouse NINJ1 primary antibody (clone 25; kind gift from Dr. Kayagaki and Dr. Dixit, Genentech, Inc.) (*19*) at 10 µg/mL in blocking solution at 4 °C overnight. The following day, cells were washed three times with PBS before the addition of secondary antibody in PBS with 1% donkey serum for 1 h at room temperature. Cy3-conjugated donkey anti-rabbit secondary antibody (Jackson ImmunoResearch #711-165-152) was used at a 1:100 dilution. Cells were washed three times with PBS and mounted in mounting medium (ProLong Diamond Antifade Mountant, Invitrogen P36965).

#### Stimulated depletion (STED)

STED microscopy was performed using a Leica TCS SP8 STED 3X microscope using HyD detectors and a 100x/1.4 oil objective. Samples labelled with Cy3-conjugated secondary antibody were excited using a white light laser at 554 nm. Emissions were time-gated 1.30-6.00 ns. A 1.5 W depletion laser at 660 nm was used at 70% maximal power. Images were acquired and deconvolved using Leica LAS X software with Lightning Deconvolution with an *xy* pixel size of 28.41 nm.

### Quantification and statistics

Statistical testing was calculated using Prism 9.0 (GraphPad Software Inc, La Jolla, CA; RRID:SCR_002798). Groups were compared using Student t test for two groups and ANOVA or Kruskal Wallis for multiple comparisons. Unless specified otherwise, all collected data was analyzed and a P value < 0.05 was considered statistically significant.

## Acknowledgments

We thank Dr. Nobuhiko Kayagaki and Dr. Vishva Dixit (Genentech Inc.) for providing the rabbit monoclonal anti-mouse NINJ1 antibody clone 25, the neutralizing anti-NINJ1 antibody clone D1, and the *Gsdmd* and *Ninj1* knockout animals. We thank Dr. John Brumell (Hospital for Sick Children) for providing pneumolysin and Janice Hicks (Hospital for Sick Children) for assistance with peritoneal macrophage isolation and culture. All imaging of the mouse and human cells was performed at the SickKids Imaging Facility at The Hospital for Sick Children.

## Author contributions

Conceptualization: JPB, AV, YW, HW, SAF, NMG, MWS, BES

Methodology: JPB, YW, AV, SAF, MWS, BES

Investigation: JPB, BM, LD, RC, AV, YW

Visualization: JPB, YW, LD, AV, BES

Funding acquisition: HW, SAF, NMG, MWS, BES

Project administration: MWS, BES

Supervision: HW, SAF, NMG, MWS, BES

Writing – original draft: JPB, BES

Writing – review & editing: JPB, AV, HW, SAF, NMG, MWS, BES

## Data and materials availability

All data are available in the main text or the supplementary materials. Requests for materials should be addressed to B.E.S.

## Supplemental Figures

**Supplemental Figure S1.**
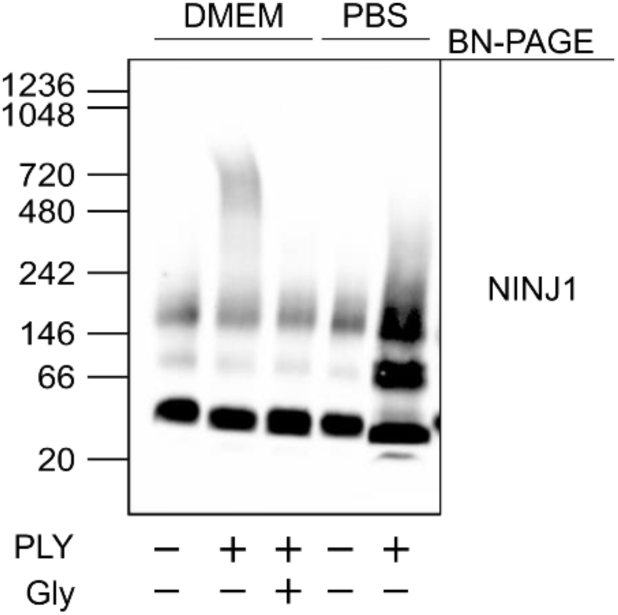
Calcium is necessary for pore-forming toxin-induced NINJ1 aggregation. Primary mouse BMDM were stimulated with pneumolysin (PLY, 0.5 mcg/mL for 15 min) in complete media (DMEM) or PBS without calcium or magnesium. Where indicated, cells were co-treated with glycine (5 mM). Western blot of NINJ1 resolved by BN-PAGE.

**Supplemental Figure S2.**
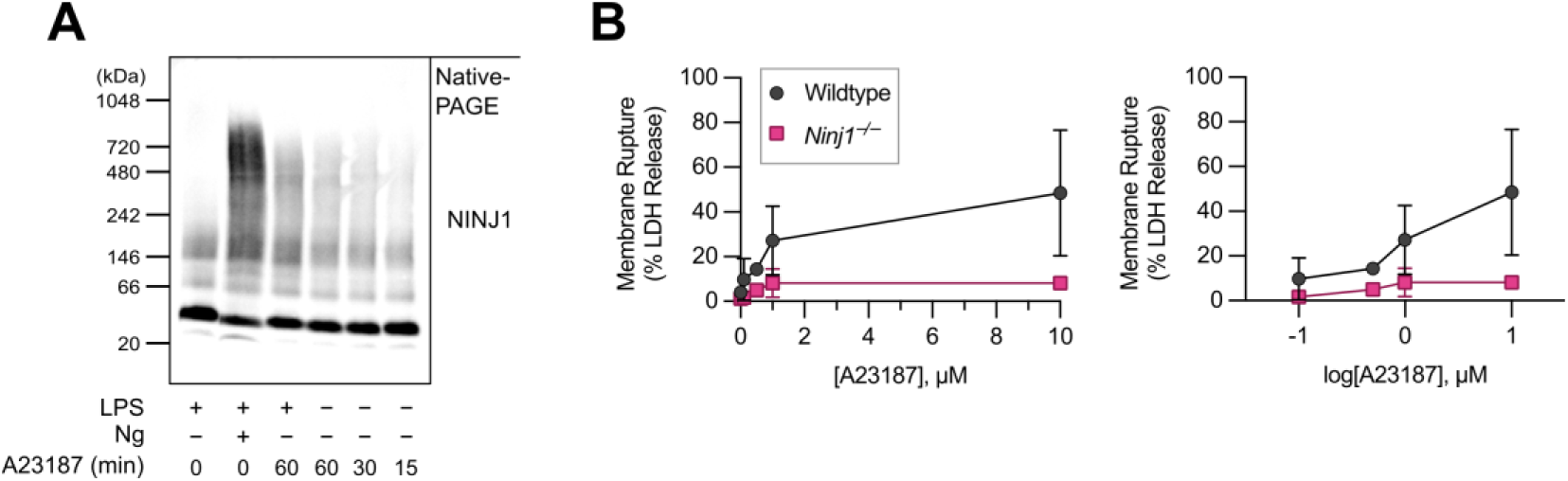
The calcium ionophore A23187 induces NINJ1 aggregation and NINJ1-dependent plasma membrane rupture. **A**, BN-PAGE analysis of LPS-primed BMDM stimulated with nigericin (20μM, 30 min) or A23187 (10 µM, 60 min), or cells stimulated with A23187 (10 µM) without LPS priming for the indicated times. **B**, LDH release from wildtype or *Ninj1*^−/−^ BMDM stimulated with A23187 at the indicated concentration for 30 min. The x-axis in each graph indicates the dose of ionomycin used to scale (left) or logarithmic scale (right).

**Supplemental Figure S3.**
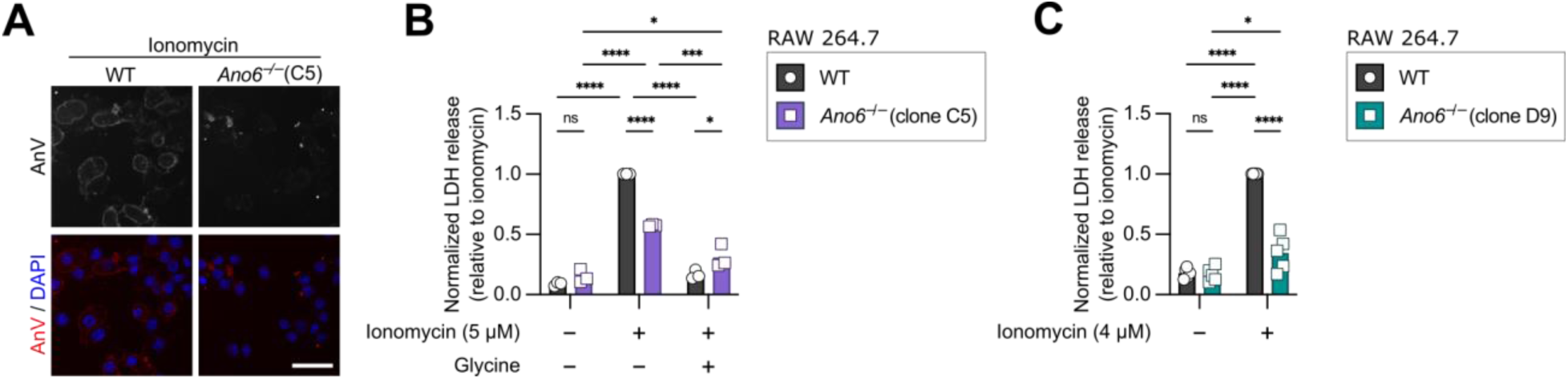
*Ano6*^−/−^ RAW 264.7 clones. Initial characterization of two *Ano6* ^−/−^ clones, named clone C5 (**A-B**) and clone D9 (**C**) generated in RAW 264.7 macrophages. Parental WT or knockout clones were treated with ionomycin for 15 min at the concentrations indicated in the figure panels. Ionomycin induces PS exposure in wildtype cells but not *Ano6* ^−/−^ clones, as indicated by annexin V labelling (**A**) PS scrambling is shown for clone D9 in Figure 2B of the main text. Ionomycin-induced LDH release is limited in *Ano6* ^−/−^ clone C5 (**B**) and clone D9 (**C**). Glycine does not offer cells from *Ano6* ^−/−^ clone C5 additional protection against ionomycin-stimulated LDH release. Data are shown normalized relative to ionomycin-treated wildtype cells. Scale bar 60 µm.

**Supplemental Figure S4.**
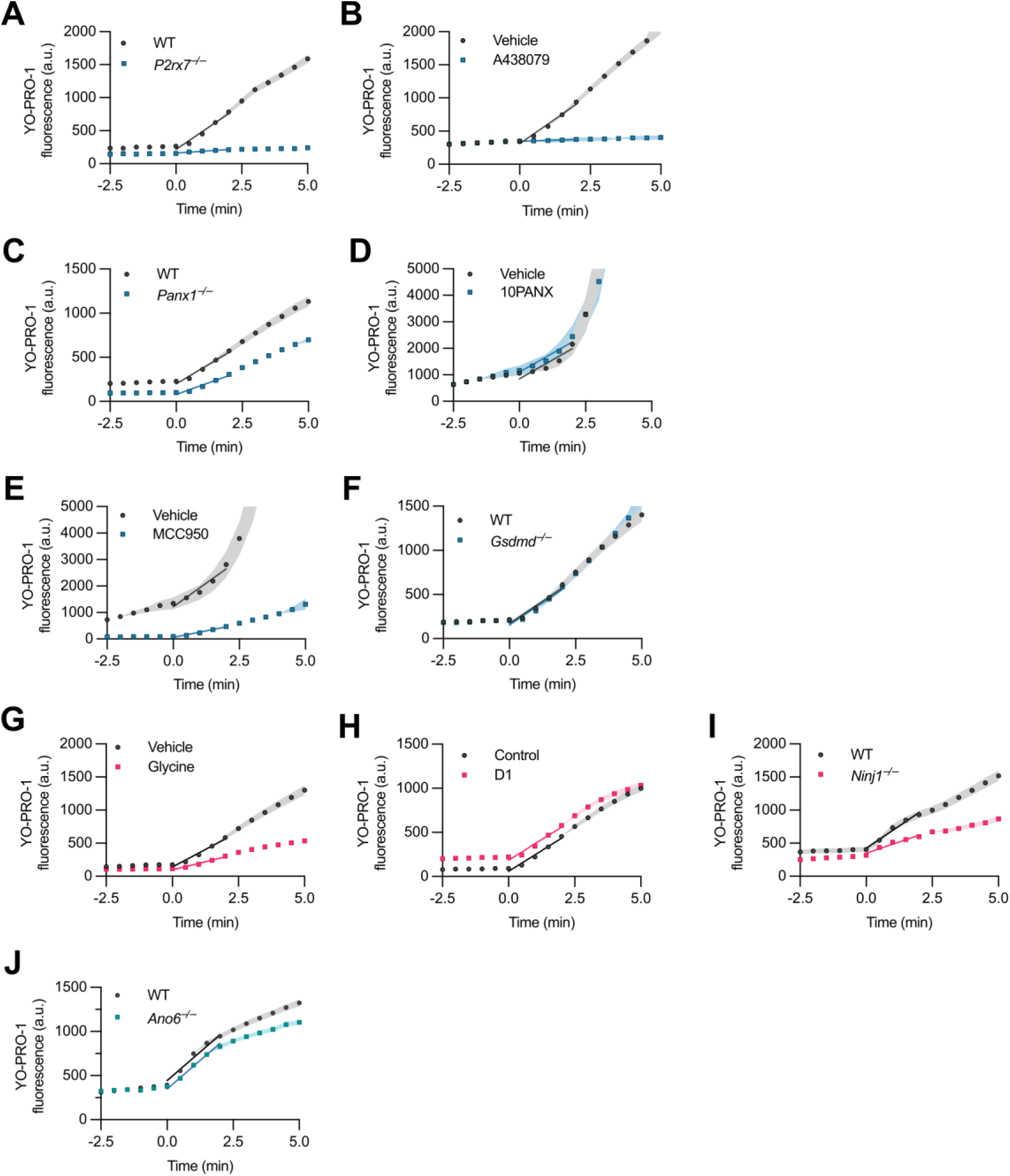
Initial linear phase of YO-PRO-1 uptake across treatment conditions and genotypes. Average YO-PRO-1 fluorescence within cells over time from t = −2.5 to 5.0 min relative to BzATP treatment for (**A**) *P2xr7* KO (**B**) A438079, (**C**) *Panx1* KO, (**D**) 10PANX, (**E**) MCC950, (**F**) *Gsdmd* KO, (**G**) glycine, (**H**) anti-NINJ1 neutralizing antibody clone D1, (**I**) *Ninj1* KO, and (**J**) *Ano6* KO.

**Supplemental Figure S5.**
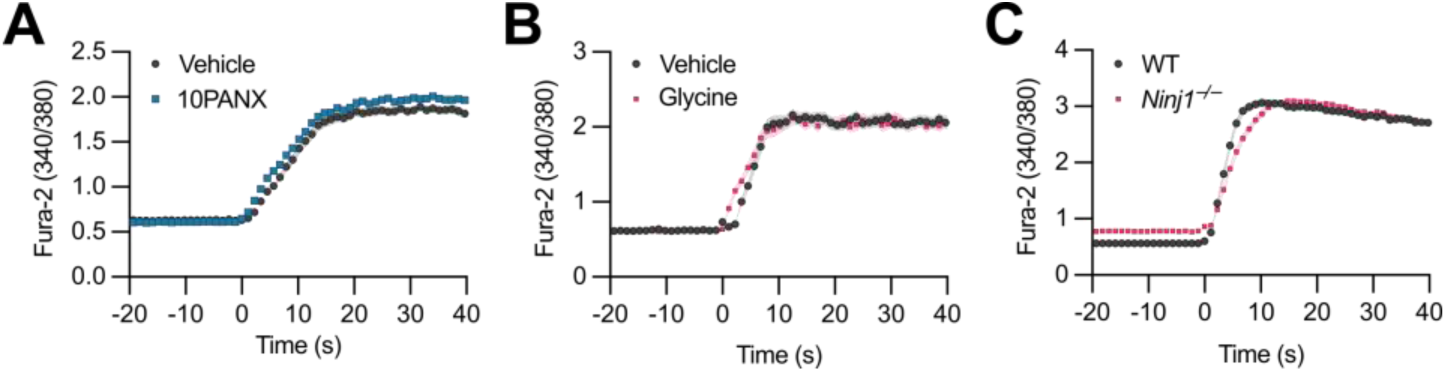
10PANX, glycine or Ninj1^-/-^ does not affect P2X7R-dependent calcium influx. **A-B**, Wildtype peritoneal macrophages were treated with BzATP (300 µM) in the presence or absence of (**A**) 10PANX or (**B**) glycine. **c**, *Ninj1* wildtype and knockout iBMDM were similarly treated with BzATP. Graphs indicate the ratio of fluorescence intensity of Fura-2 excited at 340 and 380 nm over time following BzATP treatment. Timepoints are shown relative to the first recorded measurement following BzATP treatment (t = 0 s). Data are indicative of 32-25 cells per treatment group from 1 biological replicate.

## References

1. S. Rühl, K. Shkarina, B. Demarco, R. Heilig, J. C. Santos, P. Broz, ESCRT-dependent membrane repair negatively regulates pyroptosis downstream of GSDMD activation. Science 362, 956–960 (2018).

2. A. K. Stringaris, J. Geisenhainer, F. Bergmann, C. Balshüsemann, U. Lee, G. Zysk, T. J. Mitchell, B. U. Keller, U. Kuhnt, J. Gerber, A. Spreer, M. Bähr, U. Michel, R. Nau, Neurotoxicity of Pneumolysin, a Major Pneumococcal Virulence Factor, Involves Calcium Influx and Depends on Activation of p38 Mitogen-Activated Protein Kinase. Neurobiology of Disease 11, 355–368 (2002).

3. Y. Oshimi, K. Oshimi, S. Miyazaki, Necrosis and apoptosis associated with distinct Ca2+ response patterns in target cells attacked by human natural killer cells. The Journal of Physiology 495, 319–329 (1996).

4. B. Tombal, S. R. Denmeade, J.-M. Gillis, J. T. Isaacs, A supramicromolar elevation of intracellular free calcium ([Ca2+]i) is consistently required to induce the execution phase of apoptosis. Cell Death Differ 9, 561–573 (2002).

5. L. Pedrera, R. A. Espiritu, U. Ros, J. Weber, A. Schmitt, J. Stroh, S. Hailfinger, S. Von Karstedt, A. J. García-Sáez, Ferroptotic pores induce Ca2+ fluxes and ESCRT-III activation to modulate cell death kinetics. Cell Death Differ 28, 1644–1657 (2021).

6. Y. Hirata, R. Cai, A. Volchuk, B. E. Steinberg, Y. Saito, A. Matsuzawa, S. Grinstein, S. A. Freeman, Lipid peroxidation increases membrane tension, Piezo1 gating, and cation permeability to execute ferroptosis. Current Biology 33, 1282–1294.e5 (2023).

7. N. Kayagaki, I. B. Stowe, B. L. Lee, K. O’Rourke, K. Anderson, S. Warming, T. Cuellar, B. Haley, M. Roose-Girma, Q. T. Phung, P. S. Liu, J. R. Lill, H. Li, J. Wu, S. Kummerfeld, J. Zhang, W. P. Lee, S. J. Snipas, G. S. Salvesen, L. X. Morris, L. Fitzgerald, Y. Zhang, E. M. Bertram, C. C. Goodnow, V. M. Dixit, Caspase-11 cleaves gasdermin D for non-canonical inflammasome signalling. Nature 526, 666–671 (2015).

8. R. A. Aglietti, A. Estevez, A. Gupta, M. G. Ramirez, P. S. Liu, N. Kayagaki, C. Ciferri, V. M. Dixit, E. C. Dueber, GsdmD p30 elicited by caspase-11 during pyroptosis forms pores in membranes. Proc Natl Acad Sci USA 113, 7858–7863 (2016).

9. W. He, H. Wan, L. Hu, P. Chen, X. Wang, Z. Huang, Z.-H. Yang, C.-Q. Zhong, J. Han, Gasdermin D is an executor of pyroptosis and required for interleukin-1β secretion. Cell Res 25, 1285–1298 (2015).

10. C. Virginio, A. MacKenzie, R. A. North, A. Surprenant, Kinetics of cell lysis, dye uptake and permeability changes in cells expressing the rat P2X _7_ receptor. The Journal of Physiology 519, 335–346 (1999).

11. F. Cevoli, B. Arnould, F. A. Peralta, T. Grutter, Untangling Macropore Formation and Current Facilitation in P2X7. IJMS 24, 10896 (2023).

12. R. E. Sorge, T. Trang, R. Dorfman, S. B. Smith, S. Beggs, J. Ritchie, J.-S. Austin, D. V. Zaykin, H. V. Meulen, M. Costigan, T. A. Herbert, M. Yarkoni-Abitbul, D. Tichauer, J. Livneh, E. Gershon, M. Zheng, K. Tan, S. L. John, G. D. Slade, J. Jordan, C. J. Woolf, G. Peltz, W. Maixner, L. Diatchenko, Z. Seltzer, M. W. Salter, J. S. Mogil, Genetically determined P2X7 receptor pore formation regulates variability in chronic pain sensitivity. Nat Med 18, 595–599 (2012).

13. T. Trang, S. Beggs, M. W. Salter, ATP receptors gate microglia signaling in neuropathic pain. Experimental Neurology 234, 354–361 (2012).

14. T. H. Steinberg, S. C. Silverstein, Extracellular ATP4-promotes cation fluxes in the J774 mouse macrophage cell line. Journal of Biological Chemistry 262, 3118–3122 (1987).

15. R. Muñoz-Planillo, P. Kuffa, G. Martínez-Colón, B. L. Smith, T. M. Rajendiran, G. Núñez, K+ Efflux Is the Common Trigger of NLRP3 Inflammasome Activation by Bacterial Toxins and Particulate Matter. Immunity 38, 1142–1153 (2013).

16. S. Mariathasan, D. S. Weiss, K. Newton, J. McBride, K. O’Rourke, M. Roose-Girma, W. P. Lee, Y. Weinrauch, D. M. Monack, V. M. Dixit, Cryopyrin activates the inflammasome in response to toxins and ATP. Nature 440, 228–232 (2006).

17. I. Wittig, H.-P. Braun, H. Schägger, Blue native PAGE. Nat Protoc 1, 418–428 (2006).

18. J. P. Borges, R. S. Sætra, A. Volchuk, M. Bugge, P. Devant, B. Sporsheim, B. R. Kilburn, C. L. Evavold, J. C. Kagan, N. M. Goldenberg, T. H. Flo, B. E. Steinberg, Glycine inhibits NINJ1 membrane clustering to suppress plasma membrane rupture in cell death. eLife 11, e78609 (2022).

19. N. Kayagaki, O. S. Kornfeld, B. L. Lee, I. B. Stowe, K. O’Rourke, Q. Li, W. Sandoval, D. Yan, J. Kang, M. Xu, J. Zhang, W. P. Lee, B. S. McKenzie, G. Ulas, J. Payandeh, M. Roose-Girma, Z. Modrusan, R. Reja, M. Sagolla, J. D. Webster, V. Cho, T. D. Andrews, L. X. Morris, L. A. Miosge, C. C. Goodnow, E. M. Bertram, V. M. Dixit, NINJ1 mediates plasma membrane rupture during lytic cell death. Nature 591, 131–136 (2021).

20. S. Bouillot, E. Reboud, P. Huber, Functional Consequences of Calcium Influx Promoted by Bacterial Pore-Forming Toxins. Toxins 10, 387 (2018).

21. N. Kayagaki, I. B. Stowe, K. Alegre, I. Deshpande, S. Wu, Z. Lin, O. S. Kornfeld, B. L. Lee, J. Zhang, J. Liu, E. Suto, W. P. Lee, K. Schneider, W. Lin, D. Seshasayee, T. Bhangale, C. Chalouni, M. C. Johnson, P. Joshi, J. Mossemann, S. Zhao, D. Ali, N. M. Goldenberg, B. A. Sayed, B. E. Steinberg, K. Newton, J. D. Webster, R. L. Kelly, V. M. Dixit, Inhibiting membrane rupture with NINJ1 antibodies limits tissue injury. Nature 618, 1072–1077 (2023).

22. M. Doktorova, J. L. Symons, I. Levental, Structural and functional consequences of reversible lipid asymmetry in living membranes. Nat Chem Biol 16, 1321–1330 (2020).

23. I. Shlomovitz, M. Speir, M. Gerlic, Flipping the dogma - phosphatidylserine in non-apoptotic cell death. Cell Commun Signal 17, 139 (2019).

24. N. M. de Vasconcelos, N. Van Opdenbosch, H. Van Gorp, E. Parthoens, M. Lamkanfi, Single-cell analysis of pyroptosis dynamics reveals conserved GSDMD-mediated subcellular events that precede plasma membrane rupture. Cell Death Differ 26, 146–161 (2019).

25. I. Vermes, C. Haanen, H. Steffens-Nakken, C. Reutellingsperger, A novel assay for apoptosis Flow cytometric detection of phosphatidylserine expression on early apoptotic cells using fluorescein labelled Annexin V. Journal of Immunological Methods 184, 39–51 (1995).

26. J. Ousingsawat, R. Schreiber, K. Kunzelmann, TMEM16F/Anoctamin 6 in Ferroptotic Cell Death. Cancers 11, 625 (2019).

27. A. L. Samson, Y. Zhang, N. D. Geoghegan, X. J. Gavin, K. A. Davies, M. J. Mlodzianoski, L. W. Whitehead, D. Frank, S. E. Garnish, C. Fitzgibbon, A. Hempel, S. N. Young, A. V. Jacobsen, W. Cawthorne, E. J. Petrie, M. C. Faux, K. Shield-Artin, N. Lalaoui, J. M. Hildebrand, J. Silke, K. L. Rogers, G. Lessene, E. D. Hawkins, J. M. Murphy, MLKL trafficking and accumulation at the plasma membrane control the kinetics and threshold for necroptosis. Nat Commun 11, 3151 (2020).

28. J. Suzuki, M. Umeda, P. J. Sims, S. Nagata, Calcium-dependent phospholipid scrambling by TMEM16F. Nature 468, 834–838 (2010).

29. T. Le, Z. Jia, S. C. Le, Y. Zhang, J. Chen, H. Yang, An inner activation gate controls TMEM16F phospholipid scrambling. Nat Commun 10, 1846 (2019).

30. J. Ousingsawat, P. Wanitchakool, R. Schreiber, K. Kunzelmann, Contribution of TMEM16F to pyroptotic cell death. Cell Death Dis 9, 300 (2018).

31. A. Surprenant, F. Rassendren, E. Kawashima, R. A. North, G. Buell, The Cytolytic P2Z Receptor for Extracellular ATP Identified as a P2X Receptor (P2X7). Science 272, 735–738 (1996).

32. B. R. Bianchi, K. J. Lynch, E. Touma, W. Niforatos, E. C. Burgard, K. M. Alexander, H. S. Park, H. Yu, R. Metzger, E. Kowaluk, M. F. Jarvis, T. Van Biesen, Pharmacological characterization of recombinant human and rat P2X receptor subtypes. European Journal of Pharmacology 376, 127–138 (1999).

33. T. Idziorek, J. Estaquier, F. De Bels, J.-C. Ameisen, YOPRO-1 permits cytofluorometric analysis of programmed cell death (apoptosis) without interfering with cell viability. Journal of Immunological Methods 185, 249–258 (1995).

34. R. C. Allsopp, S. Dayl, A. Bin Dayel, R. Schmid, R. J. Evans, Mapping the Allosteric Action of Antagonists A740003 and A438079 Reveals a Role for the Left Flipper in Ligand Sensitivity at P2X7 Receptors. Mol Pharmacol 93, 553–562 (2018).

35. M. Salter, J. Hicks, ATP-evoked increases in intracellular calcium in neurons and glia from the dorsal spinal cord. J. Neurosci. 14, 1563–1575 (1994).

36. S. R. Fam, C. J. Gallagher, M. W. Salter, P2Y _1_ Purinoceptor-Mediated Ca ^2+^ Signaling and Ca ^2+^ Wave Propagation in Dorsal Spinal Cord Astrocytes. J. Neurosci. 20, 2800–2808 (2000).

37. L. Erb, K. D. Lustig, D. M. Sullivan, J. T. Turner, G. A. Weisman, Functional expression and photoaffinity labeling of a cloned P2U purinergic receptor. Proc. Natl. Acad. Sci. U.S.A. 90, 10449–10453 (1993).

38. S. Hashioka, Y. F. Wang, J. P. Little, H. B. Choi, A. Klegeris, P. L. McGeer, J. G. McLarnon, Purinergic responses of calcium-dependent signaling pathways in cultured adult human astrocytes. BMC Neurosci 15, 18 (2014).

39. T. Murakami, J. Ockinger, J. Yu, V. Byles, A. McColl, A. M. Hofer, T. Horng, Critical role for calcium mobilization in activation of the NLRP3 inflammasome. Proc. Natl. Acad. Sci. U.S.A. 109, 11282–11287 (2012).

40. G.-S. Lee, N. Subramanian, A. I. Kim, I. Aksentijevich, R. Goldbach-Mansky, D. B. Sacks, R. N. Germain, D. L. Kastner, J. J. Chae, The calcium-sensing receptor regulates the NLRP3 inflammasome through Ca2+ and cAMP. Nature 492, 123–127 (2012).

41. J. Wu, A. Raman, N. J. Coffey, X. Sheng, J. Wahba, M. J. Seasock, Z. Ma, P. Beckerman, D. Laczkó, M. B. Palmer, J. B. Kopp, J. J. Kuo, S. S. Pullen, C. M. Boustany-Kari, A. Linkermann, K. Susztak, The key role of NLRP3 and STING in APOL1-associated podocytopathy. Journal of Clinical Investigation 131, e136329 (2021).

42. K. Triantafilou, S. Kar, F. J. M. Van Kuppeveld, M. Triantafilou, Rhinovirus-Induced Calcium Flux Triggers NLRP3 and NLRC5 Activation in Bronchial Cells. Am J Respir Cell Mol Biol 49, 923–934 (2013).

43. R. C. Coll, A. A. B. Robertson, J. J. Chae, S. C. Higgins, R. Muñoz-Planillo, M. C. Inserra, I. Vetter, L. S. Dungan, B. G. Monks, A. Stutz, D. E. Croker, M. S. Butler, M. Haneklaus, C. E. Sutton, G. Núñez, E. Latz, D. L. Kastner, K. H. G. Mills, S. L. Masters, K. Schroder, M. A. Cooper, L. A. J. O’Neill, A small-molecule inhibitor of the NLRP3 inflammasome for the treatment of inflammatory diseases. Nat Med 21, 248–255 (2015).

44. K. Tsuchiya, S. Hosojima, H. Hara, H. Kushiyama, M. R. Mahib, T. Kinoshita, T. Suda, Gasdermin D mediates the maturation and release of IL-1α downstream of inflammasomes. Cell Reports 34, 108887 (2021).

45. A. B. Santa Cruz Garcia, K. P. Schnur, A. B. Malik, G. C. H. Mo, Gasdermin D pores are dynamically regulated by local phosphoinositide circuitry. Nat Commun 13, 52 (2022).

46. S. Ramos, E. Hartenian, J. C. Santos, P. Walch, P. Broz, NINJ1 induces plasma membrane rupture and release of damage-associated molecular pattern molecules during ferroptosis. EMBO J 43, 1164–1186 (2024).

47. N. Kayagaki, O. S. Kornfeld, B. L. Lee, I. B. Stowe, K. O’Rourke, Q. Li, W. Sandoval, D. Yan, J. Kang, M. Xu, J. Zhang, W. P. Lee, B. S. McKenzie, G. Ulas, J. Payandeh, M. Roose-Girma, Z. Modrusan, R. Reja, M. Sagolla, J. D. Webster, V. Cho, T. D. Andrews, L. X. Morris, L. A. Miosge, C. C. Goodnow, E. M. Bertram, V. M. Dixit, NINJ1 mediates plasma membrane rupture during lytic cell death. Nature 591, 131–136 (2021).

48. Y. Dondelinger, D. Priem, J. Huyghe, T. Delanghe, P. Vandenabeele, M. J. M. Bertrand, NINJ1 is activated by cell swelling to regulate plasma membrane permeabilization during regulated necrosis. Cell Death Dis 14, 755 (2023).

49. D. B. Light, A. J. Attwood, C. Siegel, N. L. Baumann, Cell swelling increases intracellular calcium in *Necturus* erythrocytes. Journal of Cell Science 116, 101–109 (2003).

50. D. Becker, J. Bereiter-Hahn, M. Jendrach, Functional interaction of the cation channel transient receptor potential vanilloid 4 (TRPV4) and actin in volume regulation. European Journal of Cell Biology 88, 141–152 (2009).

51. S. Gatidis, O. Borst, M. Föller, F. Lang, Effect of osmotic shock and urea on phosphatidylserine scrambling in thrombocyte cell membranes. American Journal of Physiology-Cell Physiology 299, C111–C118 (2010).

52. M. Degen, J. C. Santos, K. Pluhackova, G. Cebrero, S. Ramos, G. Jankevicius, E. Hartenian, U. Guillerm, S. A. Mari, B. Kohl, D. J. Müller, P. Schanda, T. Maier, C. Perez, C. Sieben, P. Broz, S. Hiller, Structural basis of NINJ1-mediated plasma membrane rupture in cell death. Nature, doi: 10.1038/s41586-023-05991-z (2023).

53. L. David, J. P. Borges, L. R. Hollingsworth, A. Volchuk, I. Jansen, E. Garlick, B. E. Steinberg, H. Wu, NINJ1 mediates plasma membrane rupture by cutting and releasing membrane disks. Cell 187, 2224–2235.e16 (2024).

54. J. M. Whitlock, H. C. Hartzell, Anoctamins/TMEM16 Proteins: Chloride Channels Flirting with Lipids and Extracellular Vesicles. Annu. Rev. Physiol. 79, 119–143 (2017).

55. E. van den Brink-van der Laan, J. Antoinette Killian, B. de Kruijff, Nonbilayer lipids affect peripheral and integral membrane proteins via changes in the lateral pressure profile. Biochimica et Biophysica Acta (BBA) - Biomembranes 1666, 275–288 (2004).

56. J. Suzuki, D. P. Denning, E. Imanishi, H. R. Horvitz, S. Nagata, Xk-Related Protein 8 and CED-8 Promote Phosphatidylserine Exposure in Apoptotic Cells. Science 341, 403–406 (2013).

57. Y. Ryoden, K. Segawa, S. Nagata, Requirement of Xk and Vps13a for the P2X7-mediated phospholipid scrambling and cell lysis in mouse T cells. Proc. Natl. Acad. Sci. U.S.A. 119, e2119286119 (2022).

58. N. Wu, V. Cernysiov, D. Davidson, H. Song, J. Tang, S. Luo, Y. Lu, J. Qian, I. E. Gyurova, S. N. Waggoner, V. Q.-H. Trinh, R. Cayrol, A. Sugiura, H. M. McBride, J.-F. Daudelin, N. Labrecque, A. Veillette, Critical Role of Lipid Scramblase TMEM16F in Phosphatidylserine Exposure and Repair of Plasma Membrane after Pore Formation. Cell Reports 30, 1129–1140.e5 (2020).

59. C. Deisl, D. W. Hilgemann, R. Syeda, M. Fine, TMEM16F and dynamins control expansive plasma membrane reservoirs. Nat Commun 12, 4990 (2021).

60. X. Yang, X. Cheng, Y. Tang, X. Qiu, Y. Wang, H. Kang, J. Wu, Z. Wang, Y. Liu, F. Chen, X. Xiao, N. Mackman, T. R. Billiar, J. Han, B. Lu, Bacterial Endotoxin Activates the Coagulation Cascade through Gasdermin D-Dependent Phosphatidylserine Exposure. Immunity 51, 983–996.e6 (2019).

61. Y.-N. Gong, C. Guy, H. Olauson, J. U. Becker, M. Yang, P. Fitzgerald, A. Linkermann, D. R. Green, ESCRT-III Acts Downstream of MLKL to Regulate Necroptotic Cell Death and Its Consequences. Cell 169, 286–300.e16 (2017).

62. J. McCullough, A. Frost, W. I. Sundquist, Structures, Functions, and Dynamics of ESCRT-III/Vps4 Membrane Remodeling and Fission Complexes. Annu. Rev. Cell Dev. Biol. 34, 85–109 (2018).

63. L. DiPeso, D. X. Ji, R. E. Vance, J. V. Price, Cell death and cell lysis are separable events during pyroptosis. Cell Death Discov. 3, 17070 (2017).

64. R. A. Le Feuvre, D. Brough, Y. Iwakura, K. Takeda, N. J. Rothwell, Priming of Macrophages with Lipopolysaccharide Potentiates P2X7-mediated Cell Death via a Caspase-1-dependent Mechanism, Independently of Cytokine Production. Journal of Biological Chemistry 277, 3210–3218 (2002).

